# gwaRs: an R shiny web application for visualizing genome-wide association studies data

**DOI:** 10.1101/2020.04.17.044784

**Authors:** Lethukuthula L Nkambule

**Affiliations:** Sydney Brenner Institute for Molecular Bioscience, Faculty of Health Sciences, University of the Witwatersrand, Johannesburg, 2193, South Africa

## Abstract

**Summary:** Although there is an exponential increase and extensive availability of genome-wide association studies data, the visualization of this data remains difficult for non-specialist users. Current software and packages for visualizing GWAS data are intended for specialists and have been developed to accomplish specific functions, favouring functionality over user experience. To facilitate this, we have developed an R shiny web application, gwaRs, that allows any general user to visualize GWAS data efficiently and effortlessly. The gwaRs web-browser interface allows users to visualize GWAS data using SNP-density, quantile-quantile, Manhattan, and Principal Component Analysis plots.

**Availability:** The gwaRs web application is publicly hosted at https://gwasviz.shinyapps.io/gwaRs/ and R source code is released under the GNU General Public License and freely available at GitHub: https://github.com/LindoNkambule/gwaRs.

## 1 Introduction

Genome-wide association studies (GWAS) involve genotyping, usually using microarray technology, hundreds of thousands to millions of genetic variants across the human genome to identify the role of common genetic variation in the manifestation of a variety of phenotypes (traits) such as common diseases, quantitative characteristics, and drug response (Vachon, 2011). As of December 2019, more than 3700 GWAS that look at the genetic contributions of single nucleotide polymorphisms (SNPs) to human conditions or human phenotypes articles had been published, which is an increase from a total of 3639 in 2018 (Mills and Rahal, 2019; Patron et al., 2019). However, with the increase in the number of GWAS carried out and published, there are very few easy-to-use software and packages available to non-specialist users for visualizing GWAS data. To facilitate this, we have developed gwaRs, a user-friendly R shiny (Chang et al., 2020) web application, for visualizing GWAS data using quantile-quantile-, Manhattan-, SNP density-, and principal component analysis (PCA) plots. The web application is free to use and does not require registration by the user.

## 2 Materials and methods

### 2.1 Input file formats

SNP density plots use the R package CMplot (Yin, 2020) and require a tab-delimited file containing three columns, with the following mandatory header labels, in no particular order: “SNP”, “CHR”, “BP”. The same file format, as in SNP density plots, is also required for Manhattan plots, but with one extra mandatory label “P”. For QQ plots, a file containing p-values with a mandatory column header label “P” is required. Association (.assoc) files, generated using PLINK, are also accepted for SNP density-, QQ-, and Manhattan plots. Currently, gwaRs only takes .evec and .eigenvec files generated using EIGENSTRAT and PLINK respectively for PCA plots.

### 2.2 Test data

To show how gwaRs can be utilized, we explored a publicly available GWAS dataset from 90 Asian HapMap individuals initially containing 228,694 SNPs and 179,493 SNPs after quality control using PLINK (Chang et al., 2015). For our analysis, 89 individuals, 45 Han Chinese from Beijing (CHB) and 44 Japanese from Tokyo (JPT), were used as one individual was removed during QC due to missing genotype data.

## 3 Results and discussion

### SNP density

We used gwaRs to look at the number of SNPs within 1MB window size in all 22 chromosomes. As shown in figure 1A, chromosome 1 has the highest number of SNPs while chromosome 21 has the least. In gwaRs, users can change the window size to their desired output.

**Fig. 1.**
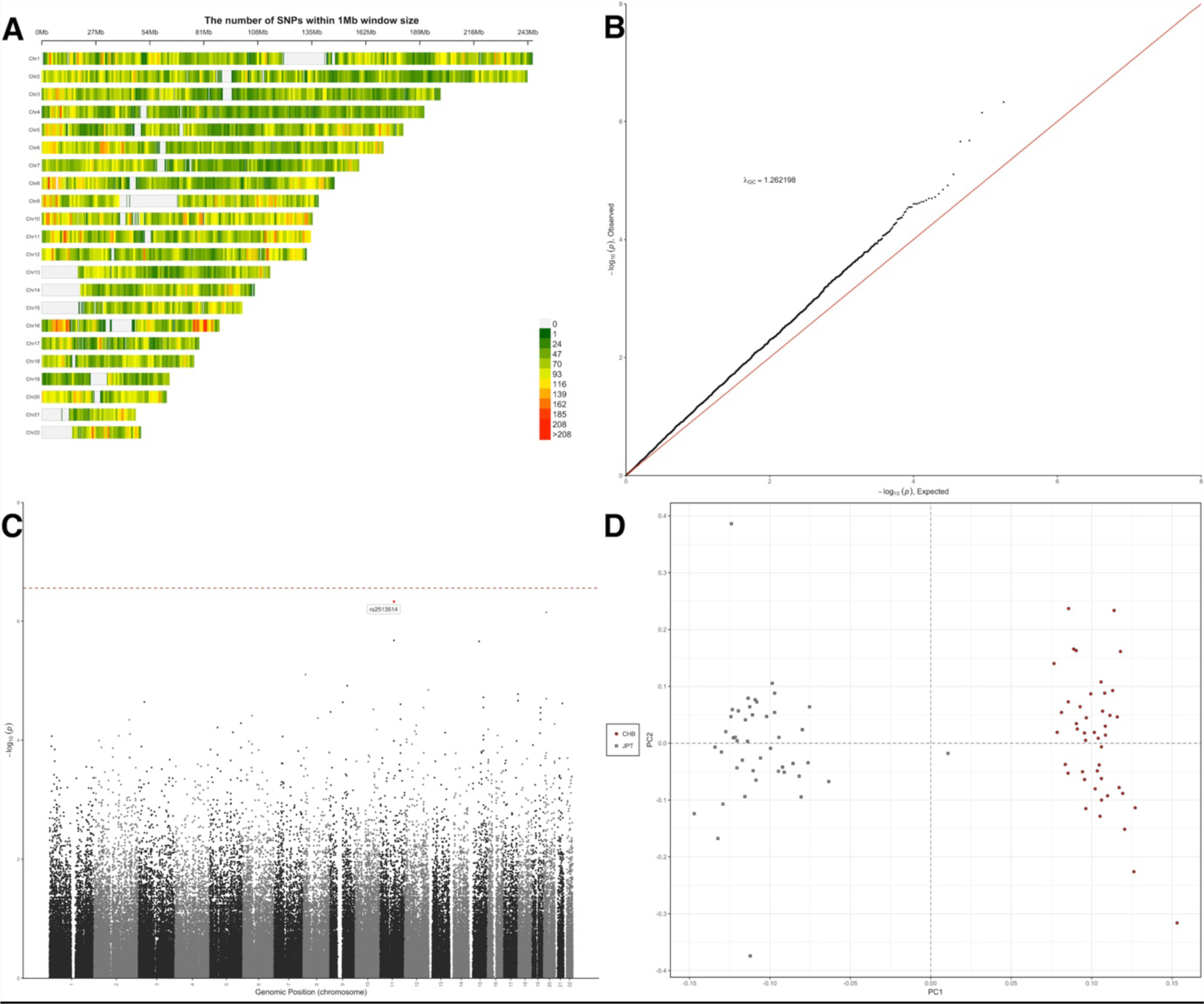
gwaRs output examples: (**A**) SNP density plot for 22 chromosomes. (**B**) QQ plot for 179,493 SNPs indicating stratification (**C**) Manhattan plot, with SNP rs2513514 annotated and highlighted in red. (**D**) PCA plot for 89 individuals.

### Quantile-quantile plots

Figure 1B shows a quantile-quantile (QQ) plot of the 179,493 SNPs. The λ_GC_ value of 1.2621 indicates that there is stratification or possible confounders such as cryptic relatedness. For association files generated using PLINK, gwaRs automatically calculates the genomic inflation factor (λ) based on median chi-squared.

### Manhattan plots

As shown in the Manhattan plot in figure 1C using the red-dotted line, no SNP reached genome-wide significance which we define as P = 2.79e^−07^ (0.05/179,493). Highlighted and annotated in the plot is SNP rs2513514 (P = 4.69e^−07^) which narrowly misses the genome-wide significance threshold. For Manhattan plots, gwaRs has two main features: (1) users can annotate SNPs by specifying a p-value threshold; (2) it allows user to generate a Manhattan plot for all the chromosomes or one specified chromosome.

### Principal Component Analysis (PCA)

The first two principal components from the filtered HapMap dataset consisting of 89 individuals are shown in figure 1D. The gwaRs PCA plot is interactive, users can click on any sample in the plot and information about the first 5 principal components, sample name and population will be shown at the bottom of the plot.

## 4 Future implementations

We are currently working on: (1) adding admixture plots; (2) switching all the current plots from static to interactive; (3) adding an option, for Manhattan plots, that allows users to annotate SNPs using rsIDs.

## Acknowledgements

We would like to thank Shaun Purcell for the test data material and Yan Holtz for allowing us to use a part of his code for Manhattan plots.

